# The Central Importance of Hub Proteins in a Disease-Gene Network Model: A New Paradigm of Chronic Myeloid Leukemia Disease Study

**DOI:** 10.1101/2021.03.21.436331

**Authors:** K M Taufiqur Rahman, Md. Fahmid Islam, Sanjib Saha, Md. Morsaline Billah

## Abstract

**Background:** The network biology of disease-gene association provides a holistic framework to decipher the intrinsic complexity of disease signaling pathways into cellular communication level. Different types of studies including large-scale genome-wide association, multifactor dimensional reduction analysis, whole genome, or exome-based sequencing strategies of diseases are striving to connect genes to diseases. Indeed, these approaches have had some accomplishments, but the cellular communication level needs a more streamlining outcome to understand the mechanistic impact of context. The higher-order combination of disease-gene interaction has a great potential to decipher the intricateness of diseases. The molecular interaction pattern of diseases at the genomic and proteomic level offers a revolutionized platform not only to understand the complexity of particular disease modules and pathways but also leading towards design novel therapeutics.

**Results:** The enrichment and topology analysis was performed by JEPETTO a plugin of Cytoscape software. We identified the chronic myeloid leukemia (CML) disease signaling pathways that appeared first in the ranking order based on XD-score among the bone, breast, and colon genes set and second at kidney and liver. This result validates the highest proximity between CML and five cancerous tissue gene set clusters. The topology analysis also supports the results while (*p<0*.*0001*) is considered to be extremely significant between CML and fives cancerous tissues genes set. Enrichment analysis identified that *abl*-gene acts as an overlapping node which is the major gene for inducing various mutations in CML. Amazingly, we identified 56 common path expansion/added genes among these five cancerous tissues which can be considered the direct cofactors of CML disease. By relative node degree, resolution, possible ligand, stoichiometry, Q-mean, and Z-score analysis we found 11 hubs proteins like SMAD3, GRB2, TP53, SMAD4, RB1, HDAC1, RAF1, ABL1, SHC1, TGFBR1, RELA which can be regarded for further drug target identification.

**Conclusions:** Our proposed network analysis reflects on the gene set interaction pattern of disease signaling pathways of humans. The integrated multidrug computational and experimental approaches boost up to improve the novel drug target approach. Besides, such a trove can yield unprecedented insights to lead to an enhanced understanding of potential application both in drug target optimization and for drug dislodging.

## Introduction

Graph theory or network biology is a complex biological model system in a wide variety of sciences like computer science, physics, interdisciplinary molecular to computational biology [1,2]. The biological network decodes the functional interaction among the biomolecules and network analysis included the potential application of drug discovery, designing protocol towards a variety of diseases, and early diagnosis of disorder for example [2, 3]. The most abundant type of biomolecules is the protein which mainly interacts with DNA, RNA, metabolites, and other protein of interest. On the other hand, protein-protein interactions (PPIs) build up a functional interface among all biological molecular pathways. Consequently, these PPIs help us to understand the inner complexity of an organism and facilitate deciphering the underlying mechanisms of diseases [4].

Network biology approaches of human diseases revolve around deciphering the interconnectedness of disease progression and identification of disease-gene signaling pathways [1]. Thus, scientists have been working for so long to achieve deeper insight to understand the molecular mechanism of disease philosophy. During the past decade, large-scale genomic data analysis has been used to achieve an intuitive understanding of disease-gene association which in turn, makes a framework to design an innovative novel therapeutic strategy [2, 3]. The functional categories of a gene fraction enrichment analysis have a better choice for gene target set analysis. The genome-wide expression profile analysis played a key role to interpret the gene sets, common biological function, chromosomal location, and regulation of genetic information [4-7]. Instead, the topology analysis can detect the distinct pattern of higher confidence disease-gene networks relationship [8, 9].

Chronic myeloid leukemia (CML) is a paradigmatic example of a malignancy characterized by karyotype abnormality and structural change of Philadelphia (Ph) chromosome through reciprocal chromosomal translocation between the long arms of 9 and 22 [10, 12]. The most prominent research suggesting the 210 fusion protein encodes *bcr-abl* hybrid gene which transcriptionally activates the pathogenesis of CML disease [13-16]. Different therapeutic strategies like stem-cell transplantation and interferon-α have been used as a quintessence treatment process, but unable to stride further for human welfare [17]. In the recent two decades, the genomic and proteomic research-driven that a single disease-causing element can act as a panacea for cancer, diabetes, and other complex diseases which are known as a magic bullet but some of them showed positive effects [18].

Now a day’ s multidrug combination therapy has emerged as a novel therapeutic approach. Indeed, multidrug interaction and combination through disease-gene interconnectedness analysis played a crucial role to develop novel therapeutics but required arduous empirical testing [19].To find out the molecular clandestine of this disease, molecular pattern [20] needs to know through comparing with other cancers like liver, kidney, bone, etc. and interactions between different pathways at the genomic and proteomic level [21, 22] should be delineated. Network biology is the primordial and fundamental approach to unfold this intricateness [23-26].

The novelty of our approach was to compare our previously published [31] five cancerous tissue proteins with KEGG database and subsequently explore the disease-gene interconnectedness by enrichment and topological signature analysis through Cytoscape 3.x plugin named JEPETTO (Java Enrichment of Pathways Extended To Topology). We have elucidated the results of cluster coefficient, degree, eigenvector, node betweenness and closeness centrality metrics to understand disease-gene correlation which will lead to identifying the functional modules and network motifs. Besides, we identified 11 hubs proteins from 56 path expansion nodes and analyzed some therapeutic target feasibility to develop drug targets against CML disease.

## METHODS

In the following section, we briefly describe how disease-gene interactions have constructed and subsequently explore the disease-gene interconnectedness by enrichment and topological signature analysis that has rendered to be helpful for further complex disease studies.

A Cytoscape 3.x plugin [33-35] named JEPETTO [27-30] offers two types of analysis; enrichment analysis and topology analysis. The genes set were submitted to the plugin for web analytical purpose. In the case of enrichment analysis, the set of genes of a protein interaction network communicates to the KEGG (Kyoto Encyclopedia of Genes and Genomes) pathways and provides a pathway ranking or tissue-specific ranking based on their XD-score. The enrichment analysis also depicts a network of interactions between input gene sets, selected pathways, and pathway expansion nodes. In the case of topology analysis, the randomly collected same size of network interactions compared to the input gene set interactions to find topological properties ordered by their topological similarities. The topology analysis also computes the topological signature of the interaction networks and compares it to the reference KEGG database. Moreover, visualize five distinct topological properties and genes mapped onto the interaction networks also a crucial finding of topology analysis.

During the enrichment analysis the parameter value set default like molecular network (default STRING), identifier format (HGNC symbol), reference database (KEGG), tissue specificity (activated), and the advanced options association threshold 1.0, coverage threshold 0.3, triangle threshold 0.1 were considered for network analysis. On the other hand, default parameters were also set for topology analysis like interaction network (large-small scale experiments), identifier format (HGNC symbol), and reference database (KEGG).

The RCSB protein data bank (http://www.rcsb.org/pdb/home/home.do) was used to identify the resolution, ligand, and stoichiometry. The Q-mean [31] and Z-score [32] predict a more valid proposition for drug target identification. The process flow (Figure 1) overview our systemic approach to develop novel drug target. The color gradient heatmap visualization was done by FunRich 2.1.2 software [47].

**Figure 1:**
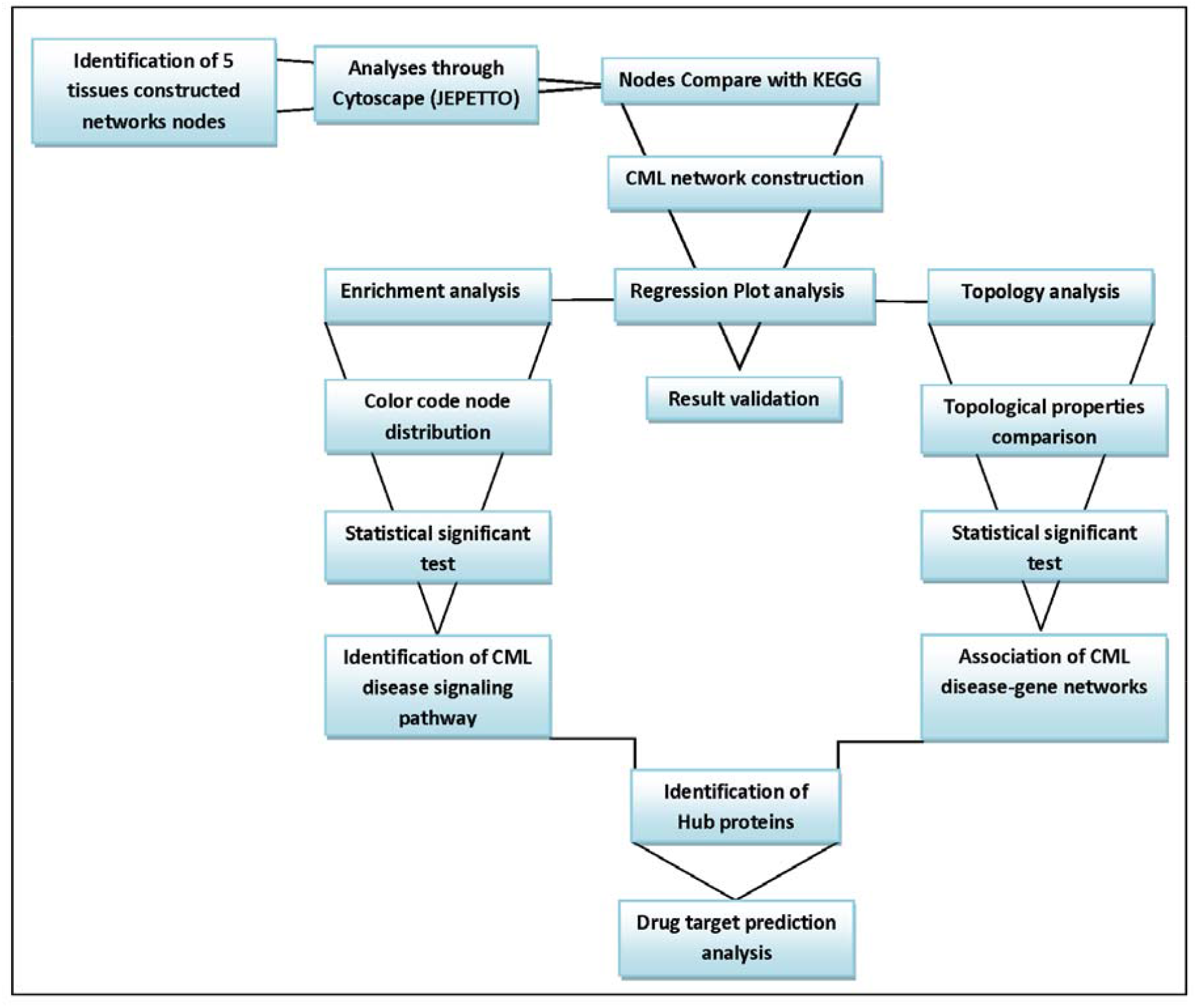
The process diagram of CML disease drug target prediction.

## RESULTS AND DISCUSSIONS

### Enrichment analysis through ranking, mapping and regression

The enrichment analysis helps us to identify the CML disease signaling pathway. The PPI networks associated genes of selected five tissues, e.g. bone, breast, colon, kidney and liver in cancer condition were used for enrichment analysis. However, the reference database KEGG compared with five cancerous tissues PPI networks constitute genes set and showed overlapping pathways or processes in ranking order. The ranking order was decided by XD-score *(significantly associated network-based score)* and a Fisher exact test q-value *(the hyper-geometric p-value after a false discovery rate multiple testing correction)* for statistical significance of each overlap through the enrichment algorithm.

**Table 1 and additional file 1** showed that in bone cancer 71% of genes mapped onto the interaction network and chronic myeloid leukemia (CML) disease signaling pathways appeared first in the ranking order based on XD-score which is more than three times higher than the significance threshold found in the regression plot (figure 2A). The XD-score determines the proximity between input genes set cluster and KEGG pathway genes cluster by their random walk [38]. It is important that, the positive value of the XD-score showed a strong association of a cluster while the negative one showed a weak association.

**Figure 2:**
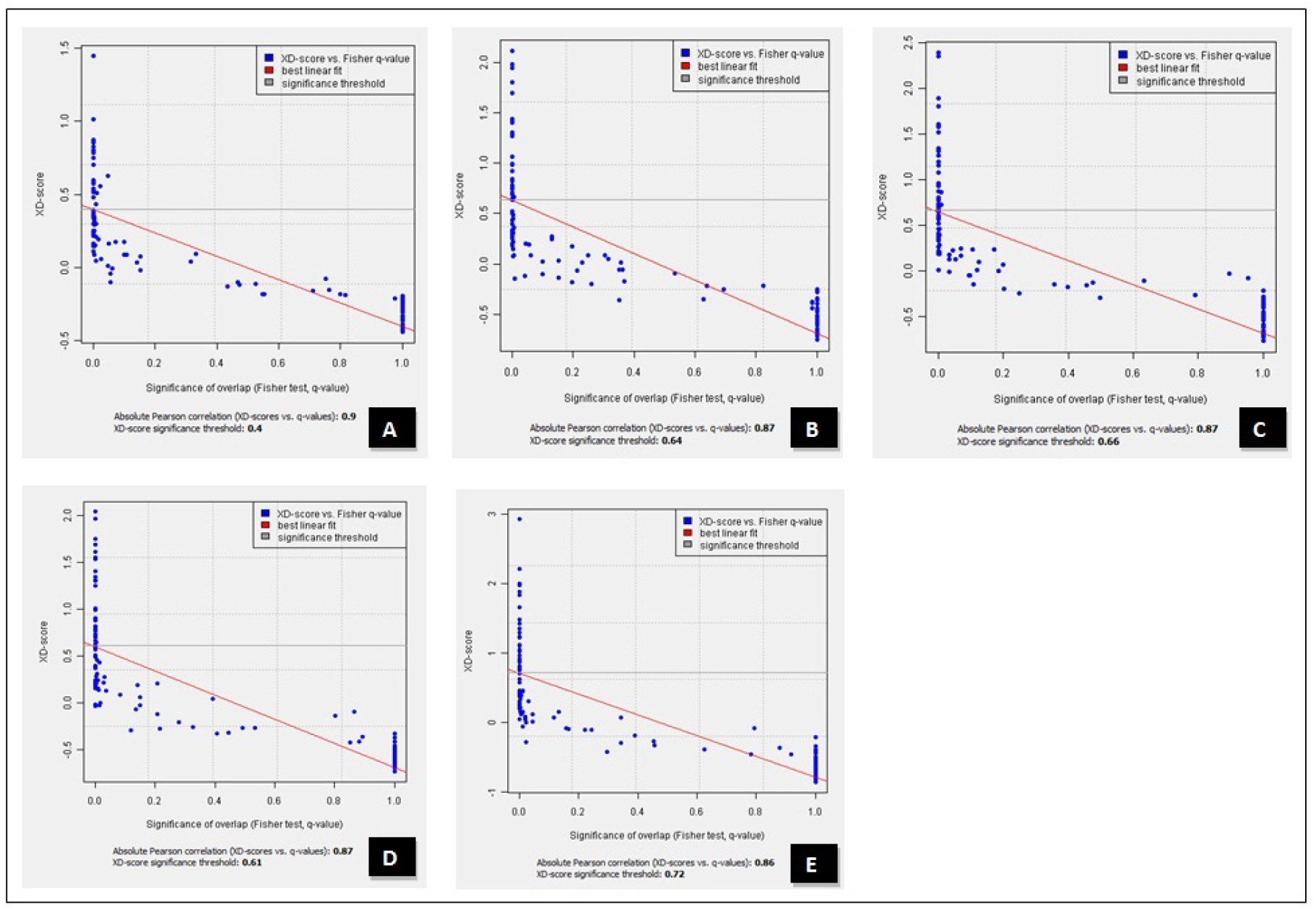
Regression plot: XD-score against the significance of overlap (Fisher test, q-value) bone (B) breast (C) colon (D) kidney (E) liver.

Intriguingly, in breast cancer, 72% of genes and in colon cancer 74% of genes mapped onto the interaction network and by comparing to the reference database CML disease signaling pathways appeared first in the ranking order. The XD-score in both cases is three times higher than the significance threshold found in the regression plot (figure 2 B, C). Instead, in kidney cancer, 74% and liver cancer 73% genes mapped, and by comparing to the reference database CML disease signaling pathways appeared third and second respectively. The XD-score in the case of the kidney approximately three and in the case of the liver more than three times higher than the significance threshold found in the regression plot (figure 2D, E). The q-value was found significant among all the cases at (*q-value<10*^*5*^) **(additional file 1)**. The regression plot indicated the XD-score vs. significance of overlap (Fisher test, q-value) which implies the best closely related disease in terms of their XD-score and q-value. The results showed that CML disease is the best fit which can provide further information to understand the biomolecular intricateness of the disease-gene network.

### Enrichment analysis of network for selected pathways

In our study, the KEGG reference database compared to the selected five cancerous tissues associated proteins and constructs a network of interaction between input gene set, selected pathway and pathway expansion nodes. In the case of the bone cancer gene set the top rank constructed network is CML based on XD-score (table 1). The constructed network showed different node information following the color code. The target set specific genes are filled with a gray color, blue for the overlap between target set and process genes, green for pathways specific genes and finally orange for the pathways expansion genes (figure 3A). Indeed, the pathway expansion genes (added nodes) are very important and we analyze this node in our results and discussion section. Then again, in the case of both breast and colon the top rank constructed network is CML (figure 3 B,C) while the kidney and liver top rank constructed network is Endometrial cancer and Notch signaling pathway respectively but the constructed signaling pathway CML ranked third and second in case of kidney and liver respectively based on their XD-score (table 1 and figure 3 D, E). So it is revealed that the cancerous protein set of five tissues is more compact in association with CML disease signaling pathway and establishing a disease-gene network to decode the complexity of molecular disease mechanism.

**Table 1.**
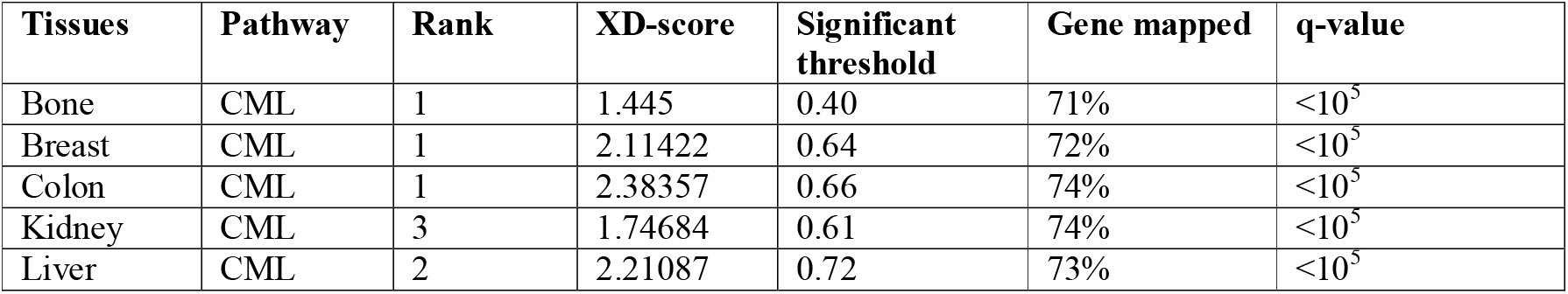
Rank based CML found in KEGG database comparing with five cancerous proteins.

**Figure 3:**
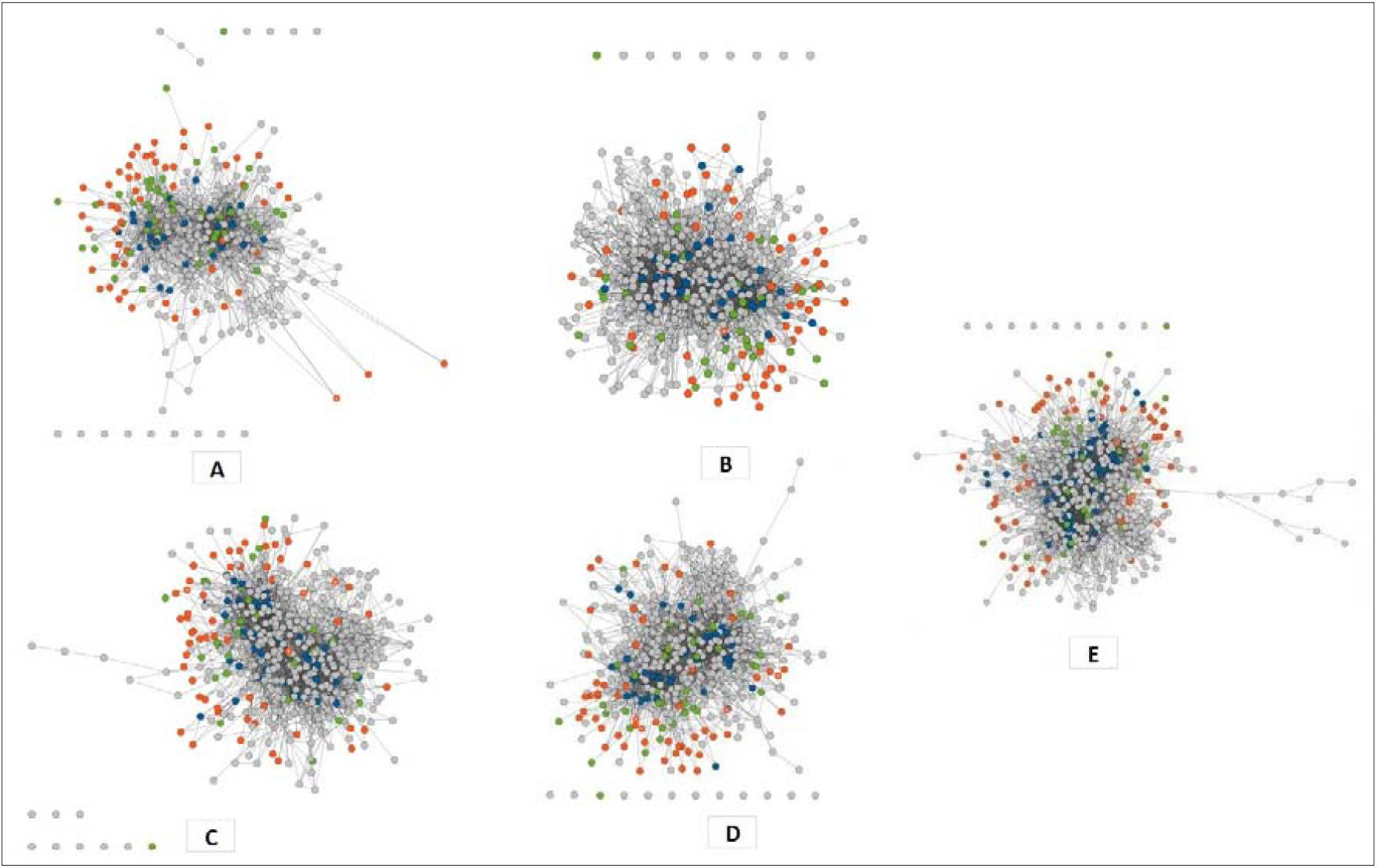
Enrichment analysis of constructing first rank CML disease signaling pathway (A) bone (B) breast (C) colon (D) kidney (E) liver.

The path expansion genes (added nodes) denote the highest closeness between two clusters. Amazingly, we found 56 common path expansion genes (added nodes) among the five tissues individual protein-protein interaction networks with CML disease signaling pathway which can be considered the direct cofactors of CML disease (additional file 2). This type of analysis can retrieve information that is not previously generated in the disease signaling mechanism. These pathway expansion nodes are crucial for further investigation to evolve a connection between cancer-forming proteins and proteins of disease signaling pathways. Inquisitively, we have found *abl-*gene as an overlapping node (additional file 3) and this gene is highly responsible for different types of mutation in CML [39, 40]. Hence, pathways protein-protein linking up can be the best interface to understand the intrinsic complexity of biological network dynamics in a more precise way.

### Topological signature analysis

Topological property analysis allowed us to reveal the close association of disease-gene networks. A randomly interacted same size of networks was used to analyze topological properties. Notably, the topological properties of a network can help to identify the anteriority of genes that are likely to be used for further functional analysis [41]. Table 2 showed that the five cancerous tissue gene sets compared to the KEGG database and provides comparable information between the uploaded gene set and 100 random simulations (mean). In the case of all the five tissues, the shortest path length is smaller than the random network that means they have closely interacted with each other. Additionally, over four times the average node degree refers that the gene set is densely connected to the target network. The target network, in this case, is CML, which carried out the same analytical results compared to the five parameters of five tissues (additional file 4).

**Table 2.** Network topological properties of 5 tissues shortest path length (SPL), node betweenness (NB), degree (D) cluster coefficient (CC), eigenvector centrality (EC)

| Tissues |  | SPL | NB | D | CC | EC |
| --- | --- | --- | --- | --- | --- | --- |
| Bone | gene set | 3.5 | 110083 | 36.85 | 0.11 | 0.08 |
| | mean | $4.13 \pm 0.04$ | $14314 \pm 4417$ | $8.03 \pm 1.33$ | $0.11 \pm 0.01$ | $0.01 \pm 0$ |
| Breast | gene set | 3.52 | 94897 | 33.68 | 0.13 | 0.08 |
| | mean | $4.13 \pm 0.04$ | $15716 \pm 5570$ | $8.47 \pm 1.34$ | $0.11 \pm 0.01$ | $0.02 \pm 0$ |
| Colon | gene set | 3.54 | 95703 | 33.84 | 0.12 | 0.08 |
| | mean | $4.12 \pm 0.02$ | $14321 \pm 3292$ | $8.23 \pm 0.76$ | $0.11 \pm 0.01$ | $0.02 \pm 0$ |
| Kidney | gene set | 3.51 | 101967 | 35.48 | 0.12 | 0.08 |
| | mean | $4.11 \pm 0.02$ | $15058 \pm 4087$ | $8.28 \pm 0.99$ | $0.11 \pm 0.01$ | $0.02 \pm 0$ |
| Liver | gene set | 3.55 | 90657 | 32.82 | 0.13 | 0.08 |
| | mean | $4.13 \pm 0.02$ | $14456 \pm 2590$ | $8.18 \pm 0.72$ | $0.11 \pm 0.01$ | $0.02 \pm 0$ |

On the other hand, the node betweenness centrality over six times higher which indicates that the presence of more central nodes (hubs) in that network. Also, the p-value result is considered to be extremely significant in between CML and fives cancerous tissues individual average node gene degree, shortest path length and node betweenness centrality (p<0.0001). Then again, the cluster coefficients p-value statistics analysis in between five tissues and CML is considered significant (p<0.0001) [additional file 5] as these signals confirm the target set are not tending to form clusters. Finally, the eigenvector centrality (EC) difference denotes the importance of the node’ s diversity of a network. In the case of our network, there are no such nodes diversity has been found.

The comparative analysis of degree against cluster coefficient; node betweenness against shortest path length of five cancerous tissues defines the closeness and high degree score among uploaded gene sets, CML disease, genetic information processing, cellular process, environmental information processing and metabolism. The plot showed that the highest degree is more than 35 and SPL is 4.6 similar among all the five tissues (figure 5). The network closeness depends on their shortest path length (SPL) score. The SPL is very similar between CML and five tissues gene set and random simulation (table 2 and Additional file 4). The regression analysis between SPL-CML (r^2^=0.367) and SPL-tissues (r^2^=0.470, overall r^2^=0.837) was found to be significant by comparative analysis (figure 4).

**Figure 4:**
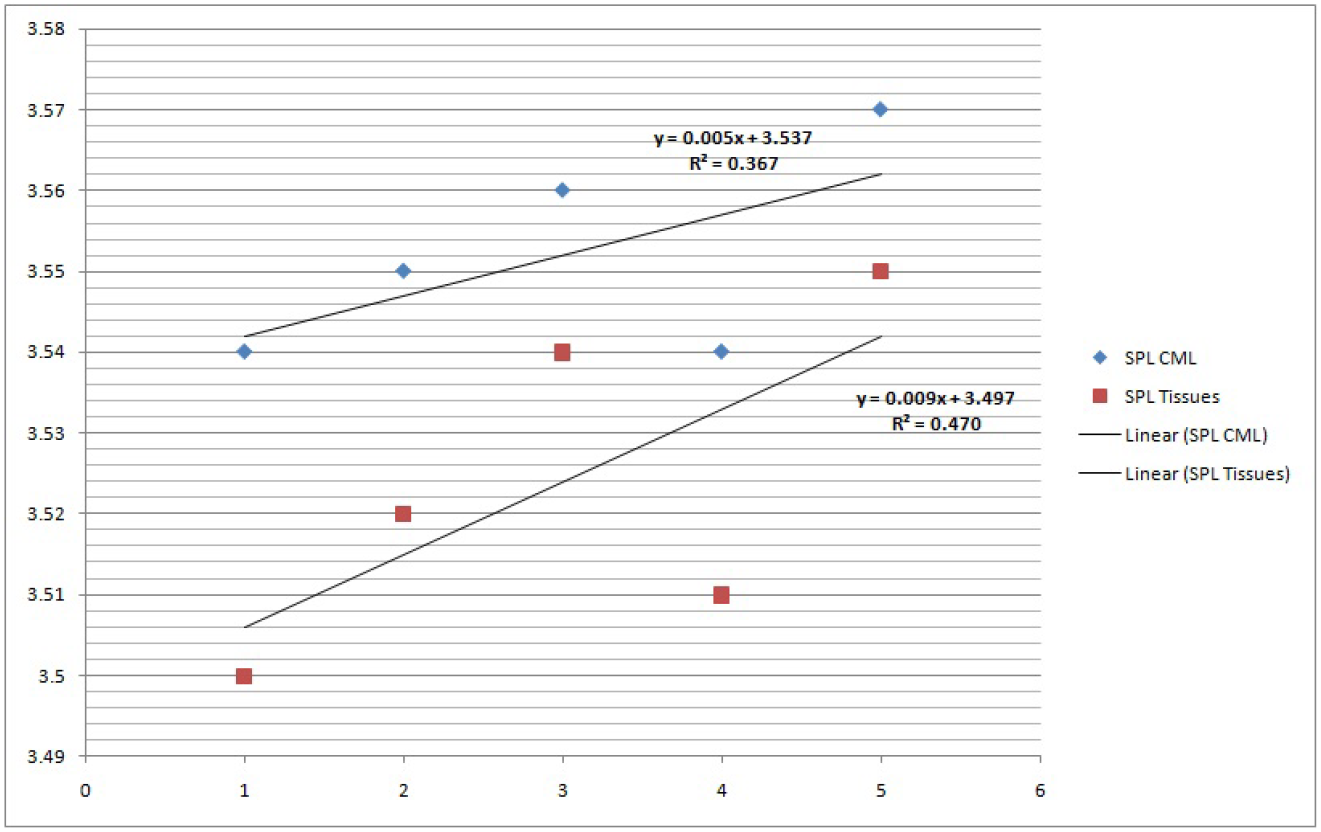
CML and five tissues regression analysis considering shortage path length.

**Figure 5:**
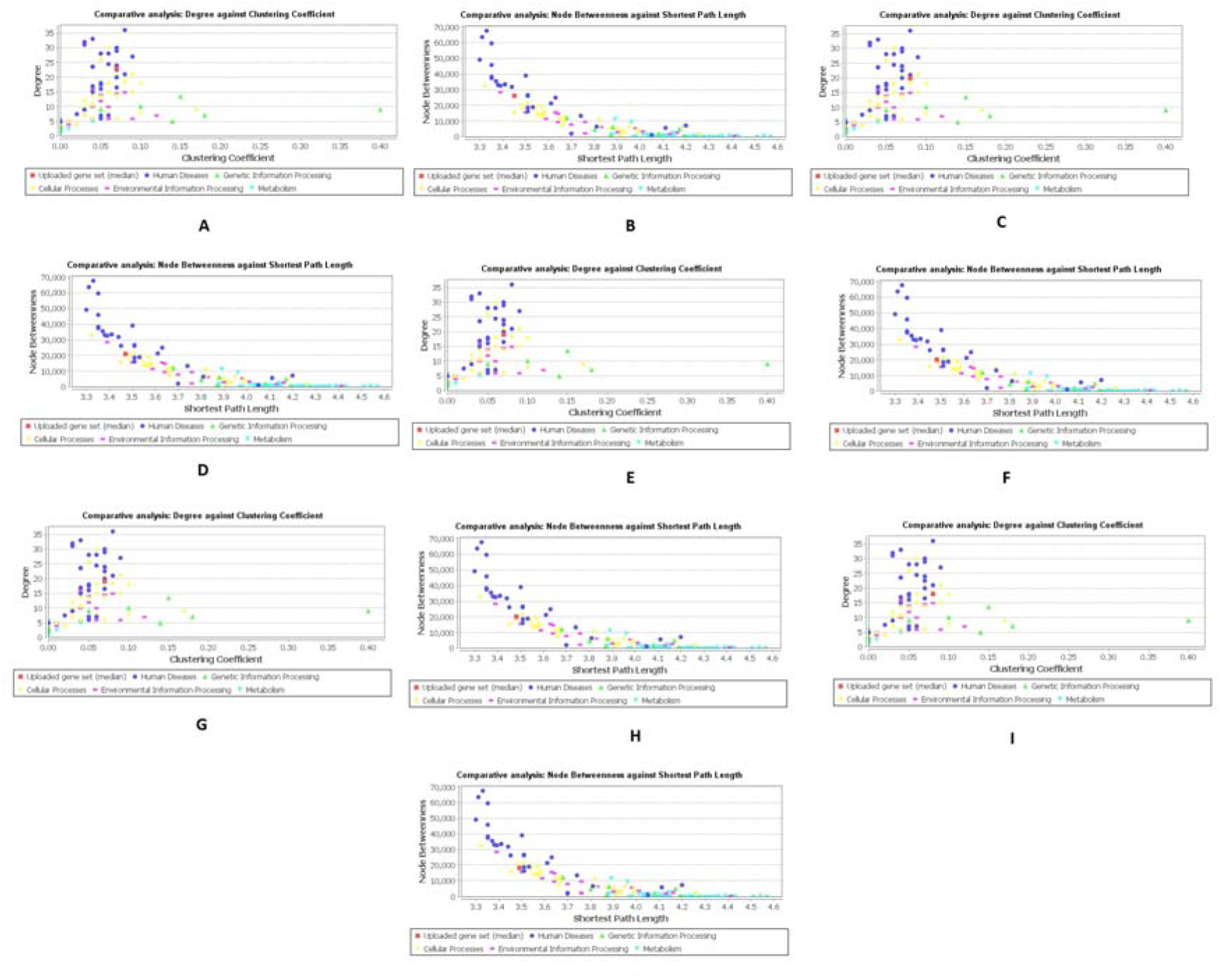
Comparative analysis: degree against cluster coefficient and node betweenness against shortage path length (A+B) bone, (C+D) breast, (E+F) colon, (G+H) kidney, (I+J) liver.

Moreover, the CML disease average node degree distribution was found highest among all human diseases in the plots. In the plots, a human disease with five distinct properties has been identified and the closeness between human disease and uploaded gene set (median) also denotes the best target disease correlation with gene sets (additional file 6).

It is noteworthy that, in the case of CML the five topological properties act remains the same in comparison to the statistics of random simulation. So the evidence showed that the constructed networks are densely connected where more hubs are present, which are the expansion/added nodes found from enrichment analysis. However, the hub proteins are dominant in a network that plays a crucial role in cellular growth and can be considered as an antimicrobial agent [42]. The knockout studies suggested that down-regulation of hub genes is possible [43] and a functional network biology approach may help to design novel therapeutics against such diseases.

### Hub proteins and drug target identification

We identified 11 highly connected nodes from 56 common path expansion nodes which clearly define the importance of nodes in biological networks. The high degree node denotes the best target for drug target identification. These nodes are mainly called the hubs or superhubs [44, 45] which are the leading subject for catalyzing vital biochemical reactions in metabolic networks [46]. Consequently, we analyze individual node’ s resolution, stoichiometry, ligand, Q-mean, and Z-score for promising drug target identification **(Table 3)**.

**Table 3:** The list of eleven nodes with the highest degree.

| Node | Resolution A<sup>0</sup> | Ligand | Stoichiometry | Q-mean | Z-score | Avg. Degree |
| --- | --- | --- | --- | --- | --- | --- |
| SMAD3 | 2.74 | 4xACY | heterotetramer | 0.679 | -0.81 | 71 |
| GRB2 | 1.6 | GOL | monomer | 0.73 | 0.20 | 65.6 |
| TP53 | 1.75 | PO4 | monomer | 0.647 | -1.31 | 54.4 |
| SMAD4 | 2.31 |  | homotetramer | 0.763 | -0.01 | 47.8 |
| RB1 | 2.4 | DHT, RAB1, SO4 | monomer | 0.847 | 0.83 | 43.2 |
| HDAC1 | 2 | SO4 | monomer | 0.757 | -0.15 | 43.6 |
| RAF1 | 2.4 | MG, PP1 | homodimer | 0.756 | 0.01 | 38.6 |
| ABL1 | 2.4 | AX1, NA, N1 | monomer | 0.774 | 0.18 | 41 |
| SHC1 | 2.41 |  | monomer | 0.733 | -0.32 | 41.8 |
| TGFBR1 | 1.7 | O85 | monomer | 0.736 | -0.41 | 38.6 |
| RELA | 2.1 | GDP, GPX, MN | monomer | 0.739 | -0.24 | 37.8 |

To be the best possible drug target, the resolution of the protein must be minimum as possible. Resolution of ∼3Å is the minimum requirement for drug design. Normally, more than 2Å is best for in silico drug design. Here, all the proteins have optimum resolution ranging from 1.6 to 2.74Å to be started working with as drug target. It is always better to start with protein-ligand complex rather than protein without any ligand because it provides the exact binding site and mode for ligands. Here, all the protein exists as protein-ligand complex except SMAD4 and SHC1. Sufficient literates are available for each protein, which can be a useful source of information of different binding sites whether single or multiple for different ligands and possible conformational changes. From the literature, we can avail the structure of binding mode and active domain, which are crucial for drug design. The correct quaternary structure of the protein is important for drug design. It is always simple and handy to use a monomeric structure if the oligomeric structure is not important for ligand binding. If the oligomeric form affects then correct oligomer should be used because it is necessary for correct folding and sometimes ligand binding site exists between monomers in the dimer. Here, SMAD3, SMAD4 and RAF1 exist as an oligomer and other proteins are simple monomers, which are simple to start with.

On the other hand, it is expected that the Q-mean score range from 0 to 1 considered for the best reliable candidates and the Z-score in maximum cases is close to 0. In our findings, all 11 proteins Q-mean and Z-score are significant.

### Heatmap correlation matrix analysis

The stem cell model of leukemia suggested that during normal hematopoiesis phage the formation of leukemia stem cells in myeloid leukemia caused different types of mutation in cells. These mutations have facilitated the development of different leukemia clones like B cell, erythrocyte, granulocyte, natural killer cell, monocyte, megakaryocyte-erythrocyte progenitor, platelet and T cell. The CD4+ and CD8+ T cells have been activated by leukemia cells secreted antigens that can control leukemia cells by deactivating the therapy-resistant leukemic stem cells [48].

In this formalism, the heatmap graphical view (figure 6) showed that the proteins correlate with the B cells, CD4 cells, CD8 cells, NK cells, monocytes, and platelets which are the major enhancer for CML disease progression. Here, GRB2 has shown the best result in comparison to the other proteins. Also, the HDAC1 and RELA have shown comparatively good results than others. In the case of GRB2 the best score in blood platelets and a very similar score to the B cells, CD8 cells, and monocytes. The HDAC1 showed the highest score in B cells and CD8 cells and similar to the rest of the cells except platelets. The third best score found in RELA and SMAD3, ABL1, SHC1, SMAD4, TP5 proteins have scored below 40 consecutively. So, we can predict that our proposed proteins of interest can be a good choice for further research.

**Figure 6:**
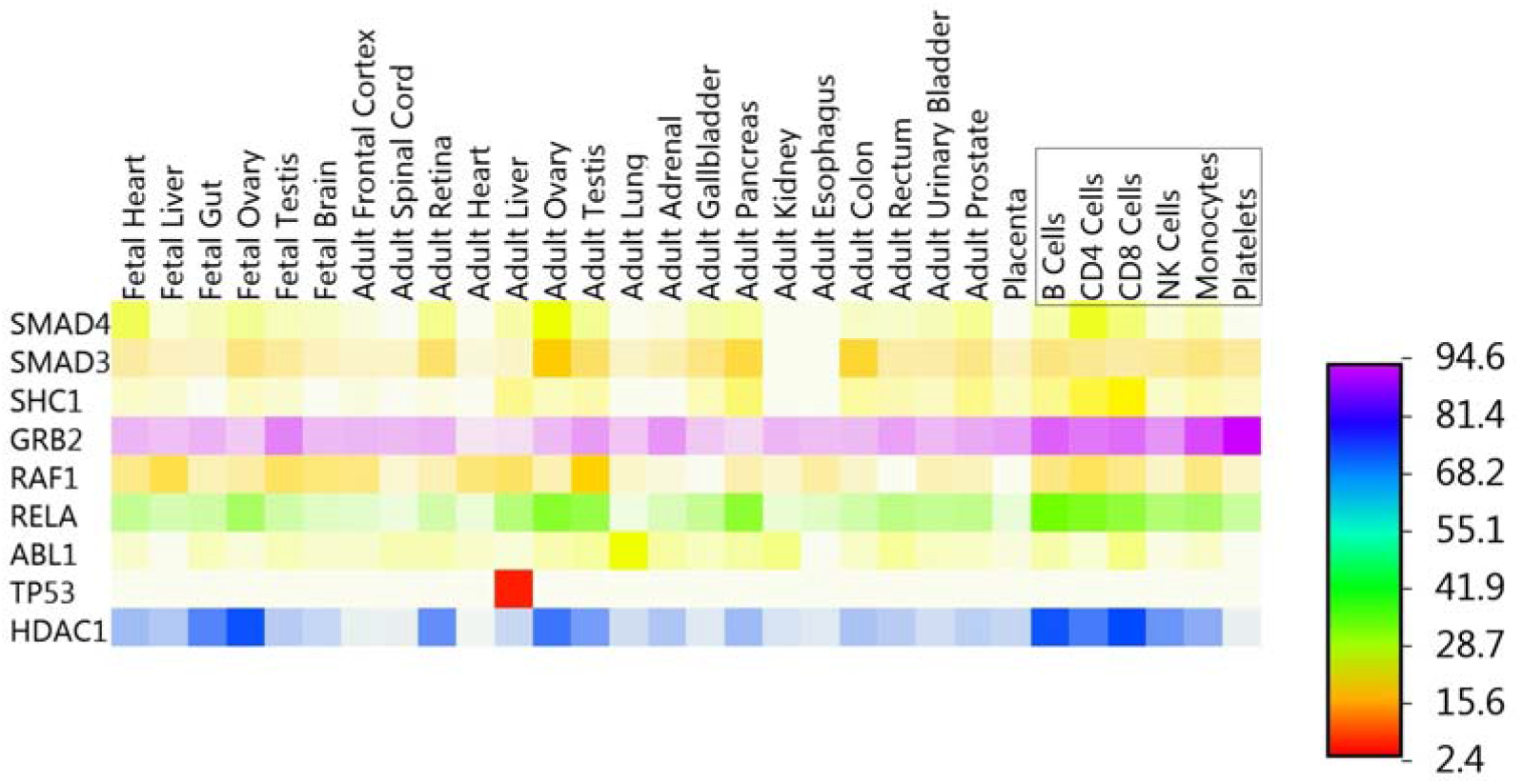
Proteins heatmap correlation matrix generated from FunRich.

## Conclusion

The crucial experiment of disease signaling pathway grasped out the transcriptome network analysis and pathway crosstalk analysis, which will be further steering the comprehensive molecular research of CML disease treatment. Our approach mainly focused on revealing the dynamic processes of CML disease in affiliation with the five cancerous tissues PPIs networks constitute genes. The analysis of PPIs network dynamics can unfold the biomolecular intricateness of the disease-gene relationship. Our study directly related to topology and enrichment analysis which play an important role to delineate network properties holistically. The hubs protein on the other hand considered the power-house of a network. The relative analysis of therapeutic target feasibility of these hubs and correlation matrix compiling to cells can be a benefit to some extent. Indeed, our theoretical approaches would be the hope for future bioinformatics-based therapeutics studies.

## Author Contributions

## Additional files

**Additional file 1:** Ranking-based disease signaling pathways found in KEGG comparing with five tissues cancerous proteins.

**Additional file 2:** Identification of 56 path expansion genes/added nodes among the five tissues.

**Additional file 3:** Identification of *abl-*overlapping nodes among the five tissues gene set.

**Additional file 4:** Network topological properties of CML considering five tissue distinct parameters.

**Additional file 5:** Significant test results considering five parameters in between five tissues and CML disease.

**Additional file 6:** Five distinct topological properties of uploaded gene set among the five tissues.

